# Frequency-Resolved Connectivity Graphs for EEG-Based Classification of Alzheimer and Frontotemporal Dementia

**DOI:** 10.64898/2026.10.16.759776

**Authors:** Mohamed Radwan, Pedro G. Lind, Anis Yazidi

**Affiliations:** Department of Computer Science, Oslo Metropolitan University, 0167 Oslo, Norway; Department of Technology, Kristiania University of Applied Sciences, 0107 Oslo, Norway; Numerical Analysis and Scientific Computing Department, Simula Research Laboratory, 0164 Oslo, Norway; Department of Informatics, University of Oslo, 0373 Oslo, Norway

**Keywords:** Graph neural networks, EEG, Alzheimer’s disease, Frontotemporal dementia, Frequency bands, Explainability

## Abstract

In this study, we build EEG based graph model that uses frequency-resolved connectivity features, with Coherence, phase locking value (PLV) and phase lag index (PLI) in five frequency bands as multi-dimensional edge features. The model is used to classify Alzheimer’s disease (AD) and frontotemporal dementia (FTD) against healthy controls (CN). This paper extends our earlier work on FTD [1] with AD vs CN as the main new task, a feature ablation for it, and a joint analysis of both tasks. With leave-one-subject-out (LOSO) validation, the model reaches an accuracy of 81.13% for AD vs CN and 76.06% for FTD vs CN, which is competitive with the baselines reported with the dataset with far fewer parameters (159 and 282 thousand versus 170.5 million). Under the random cross-validation setting used for the DICE-net model, our model has a comparable accuracy for FTD and a lower accuracy for AD. Node and edge saliency maps highlight occipital and temporal electrodes, the Alpha band for node features and the Theta band for edge features, which is consistent with earlier EEG studies. In an ablation on the AD task, using all 15 edge features performed better than using a single connectivity measure or edge features averaged over the bands.

## 1 Introduction

Alzheimer’s disease (AD) is a neurodegenerative disorder that causes cognitive decline, memory loss and changes in behavior, and its pathogenic mechanism is still poorly understood [2]. Frontotemporal dementia (FTD) is another neurodegenerative disorder, which mainly affects the frontal and temporal lobes. One route to understanding these diseases and diagnosing them is to find reliable biomarkers. Studying functional connectivity between brain regions can be helpful in the biomarker discovery. Changes in connectivity patterns can appear before clinical symptoms emerge and before structural changes are visible [3], and they may be detectable before the accumulation of beta-amyloid plaques and tau proteins [4]. Studying connectivity separately in different frequency bands can give more detail about the pathology.

Electroencephalography (EEG) is a useful tool for studying brain connectivity because it is non-invasive and has a high temporal resolution. Conventional statistical methods have been used to extract EEG features that separate patients from healthy individuals, and deep learning methods are increasingly used for the same purpose. Deep learning models are often seen as black boxes, which makes it hard to understand why a model makes a prediction. This is a concern in clinical settings, so there is a growing interest in interpreting what these models learn, in the hope of identifying robust biomarkers.

Graph neural networks (GNNs) represent EEG electrodes as nodes and the relations between electrodes as edges. In many GNN studies the adjacency matrix is built from the spatial distance between electrodes, with the assumption that close electrodes are more strongly connected. However, spatial closeness does not guarantee functional connectivity, and binary adjacency matrices discard the strength of the connection. Using functional connectivity to build a weighted adjacency matrix avoids these problems.

Several studies have used graph-based models for EEG. The authors of [5] used graph attention networks for general EEG classification without an interpretation. The authors of [6] introduced a dynamic graph CNN (DGCNN) that learns the adjacency matrix during training and applied it to emotion recognition. The authors of [7] developed a graph input layer attention convolutional network that uses Pearson correlation as the adjacency matrix, with activity, mobility, complexity and power spectral density as node features, on a depression dataset of 60 subjects. They used the weight matrix of the input layer to find important edges and nodes, but did not study frequency bands. A Chebyshev-polynomial GNN with explanations was developed in [8] to detect Parkinson’s disease from EEG. It quantifies the importance of nodes and edges to identify abnormal connectivity, and it uses microstates to study brain dynamics, but it does not use different frequency bands.

Interpretation has also been studied for CNN-based EEG models. The authors of [9] classified mild cognitive impairment (MCI) in two EEG datasets using four frequency bands separately, and found that the Beta band (15–30 Hz) gave the best performance. The authors of [10] classified abnormal EEG in the TUH dataset [11] using five frequency bands, and found the Delta band (0.5–4 Hz) to be the most important. That study uses a large dataset but a general normal and abnormal labels with limited conclusions about specific diseases. Frequency content can also be related to higher brain functions such as memory and learning, which can help both explainability and performance [12].

The work most related to ours is [13], which used GNNs for Alzheimer’s disease and epilepsy and studied feature importance for Alzheimer’s disease on the dataset of [14]. The authors report the importance of the Alpha band and of occipital (O1) and prefrontal (Fp1) regions, using PLV, Pearson correlation and cross spectral density to build the adjacency matrices. Their work focuses more on model development, while our focus is on the interpretation of the model, the ablation of the input features, and the salient features that the model uses.

### Relation to our earlier work

This paper extends our earlier work on FTD vs CN [1]. The main task here is AD vs CN, which is new, and we add a feature ablation and node and edge saliency maps for it. The FTD vs CN task from the earlier paper is included to compare the two diseases with the same model, and its results are the same as in the earlier paper.

Our contributions are:

- A frequency-resolved multi-edge GNN that uses Coherence, PLV and PLI in five frequency bands as edge features, evaluated for AD vs CN and, from our earlier work, for FTD vs CN, with leave-one-subject-out validation.
- An ablation of the edge features on the AD vs CN task, comparing each connectivity measure alone and edge features averaged over the frequency bands with the full set of 15 edge features.
- An interpretation of the model with saliency maps for nodes, edges and frequency bands in both tasks, compared with findings from earlier EEG studies.

The rest of the paper is organized as follows. Section 2 describes the data, the model, the features and the experiments. Section 3 presents the classification results, the saliency maps and the ablation. Section 4 discusses the results, and Section 5 concludes and lists the limitations.

## 2 Data and Methods

### 2.1 Data

We use the EEG dataset of [15], which contains resting-state, eyes-closed recordings from 88 subjects (36 with Alzheimer’s disease (AD), 23 with frontotemporal dementia (FTD) and 29 healthy controls (CN)) collected at the AHEPA University Hospital in Thessaloniki, Greece. The AD vs CN task therefore uses 65 subjects and the FTD vs CN task uses 52. The recordings have 19 scalp electrodes (plus 2 reference electrodes) and a sampling rate of 500 Hz. We use the preprocessed version provided by the dataset authors, in which the signals were band-pass filtered between 0.5 and 45 Hz, re-referenced to A1–A2 and cleaned of artifacts with Artifact Subspace Reconstruction (ASR) and Independent Component Analysis (ICA) following the preprocessed dataset from [15].

### 2.2 Graph Neural Network

As said, we are proposing a graph model in this study. For each time window, a graph is built with the 19 electrodes as nodes, the node features of Section 2.3, and 15 edge features for each pair of electrodes which are three connectivity measures in five frequency bands (Section 2.4). The graph is passed to Transformer Convolution layers [16], which use attention message propagation and also use the edge features in the attention. Here, we use five attention heads. The outputs of the attention heads are concatenated and followed by a ReLU activation and batch normalization (Fig. 1). After the last layer, global mean pooling over the nodes gives a graph embedding, which is passed to a fully connected layer. The model has two layers for AD vs CN and three layers for FTD vs CN, so the numbers of trainable parameters are 159,234 and 281,954.

**Figure 1:**
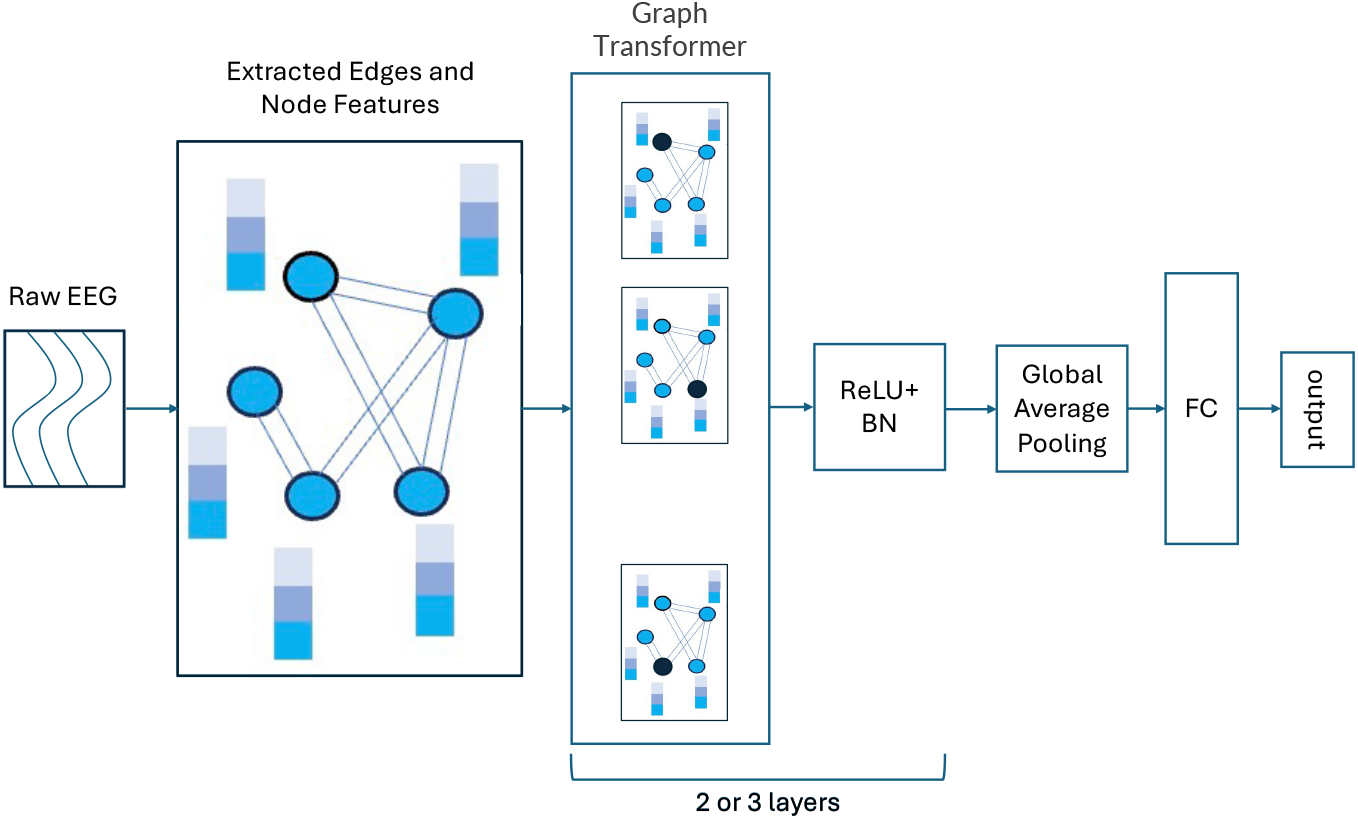
Model architecture. ReLU is the rectified linear unit, BN is batch normalization and FC is a fully connected layer. Edge and node features are calculated from the EEG data, which gives a graph for each sample. The Transformer Convolution layer followed by ReLU and BN is repeated twice for AD vs CN and three times for FTD vs CN, and the node embeddings are pooled by global average pooling and passed to an FC layer. The number of attention heads is five.

### 2.3 Frequency Bands, Windows and Node Features

The signals are decomposed with Morlet wavelets into five frequency bands, Delta (1–4 Hz), Theta (4–8 Hz), Alpha (8–13 Hz), Beta (13–30 Hz) and Gamma (30–45 Hz), and the node and edge features are calculated separately for each band. The recordings are divided into windows of 30 seconds with an overlap of 15 seconds, which is the window size used in [17] on this dataset. Each window is one sample, so the classification is window-based. The node features are hand-crafted: differential entropy, spectral entropy, permutation entropy, entropy power, approximate entropy, Hjorth mobility and complexity, fractal density and power spectral density, together with the mean, standard deviation and kurtosis of the signal.

### 2.4 Functional Connectivity

Spectral coherence, the phase locking value (PLV) and the phase lag index (PLI) quantify the relation between pairs of electrodes in each of the five bands. All three are symmetric, so the graphs are undirected, and the 15 edge features form a tensor *A ∈* ℝ^19*×*19*×*15^.

#### Spectral coherence

The magnitude-squared coherence is estimated with Welch’s method,

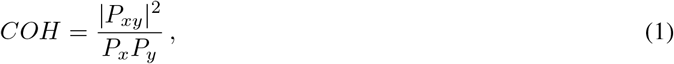

where *P*_*x*_ and *P*_*y*_ are the power spectral density estimates of the signals *x* and *y*, and *P*_*xy*_ is their cross spectral density.

#### Phase locking value (PLV)

The PLV [18] measures how consistent the phase difference between two signals is over time, ignoring the amplitudes,

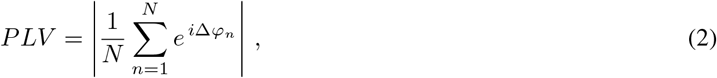

where Δ*φ*_*n*_ is the phase difference at time point *n*, the instantaneous phases are obtained from the Hilbert transform, and *N* is the number of time points. The PLV takes values between 0 (no phase synchrony) and 1 (constant phase difference).

#### Phase lag index (PLI)

The PLI [19] is less affected by a common source or volume conduction, which can inflate the PLV, because it ignores phase differences centered around 0 or *π*. It measures the asymmetry of the distribution of phase differences,

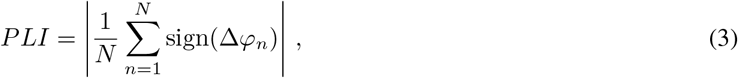

and takes values between 0 (symmetric distribution around 0 or *π*) and 1.

### 2.5 Explainable AI Methods

We use saliency maps, the gradient of the model output with respect to the input, to find which nodes and edges the model relies on. The absolute value of the gradient is taken as the feature importance [20], and the values are averaged over the data to obtain a global importance.

### 2.6 Experiments

The data are split with leave-one-subject-out (LOSO) validation, as in the baselines of [15]. In each fold, one subject is held out and the model is trained on the remaining subjects. Because the classification is window-based, the predictions for the windows of the held-out subject are combined by majority voting. We report accuracy, sensitivity, specificity and F1 score, averaged over the folds. With 65 subjects for AD vs CN and 52 for FTD vs CN, one subject corresponds to about 1.5 and 1.9 percentage points of accuracy, respectively. The two binary tasks are AD vs CN, which is the main task of this paper, and FTD vs CN, which comes from our earlier work.

The model is trained with the Adam optimizer, a learning rate of 0.00001, a batch size of 100, a dropout of 0.5, mean pooling and 35 epochs (Table 1). The small learning rate was used to avoid instability, and beyond 35 epochs the training showed signs of overfitting. The source code is available at https://github.com/mhmdrdwn/ftd_graph.

**Table 1:** Selected hyperparameters. The number of layers was two for AD vs CN and three for FTD vs CN.

| Parameter | Search space | Selected value |
| --- | --- | --- |
| Learning rate | $10^{-6} - 10^{-2}$ | $10^{-5}$ |
| Number of epochs | 0 – 200 | 35 |
| Batch size | 16 – 256 | 100 |
| Number of layers | 1 – 5 | 2 (AD), 3 (FTD) |
| Dropout | 0.1 – 0.5 | 0.5 |
| Pooling | Mean, Max, Add | Mean |

## 3 Results

### 3.1 Classification Performance

Two comparisons are made, and they use different evaluation settings. Tables 2 and 3 compare our model with the classical baselines reported with the dataset [15] using LOSO validation. Tables 4 and 5 compare it with the deep learning models of [17], whose results come from random cross-validation (ten runs, with six subjects in the validation sets), so our model was also evaluated in that setting for these two tables. The numbers of the two groups of tables are therefore not directly comparable.

**Table 2:** Comparison between our model and the baselines of [15] for Alzheimer’s disease (AD) vs healthy controls (CN). The results are the weighted average of the LOSO cross-validation performances. The best value in each column is in bold.

| AD vs CN | Accuracy | Sensitivity | Specificity | F1 Score |
| --- | --- | --- | --- | --- |
| kNN | 69.67% | 75.98% | 72.81% | 72.81% |
| SVM | 71.89% | 75.98% | 73.74% | 73.74% |
| MLP | 73.00% | 74.63% | 74.82% | 74.82% |
| LightGBM | 76.43% | 76.01% | 76.16% | 76.12% |
| Random Forest | 77.01% | 78.32% | **80.94%** | 75.31% |
| Our Model | **81.13%** | **83.05%** | 78.85% | **82.67%** |

**Table 3:** Comparison between our model and the baselines of [15] for frontotemporal dementia (FTD) vs healthy controls (CN). The results are the weighted average of the LOSO cross-validation performances. The best value in each column is in bold.

| FTD vs CN | Accuracy | Sensitivity | Specificity | F1 Score |
| --- | --- | --- | --- | --- |
| kNN | 67.34% | 59.67% | 76.13% | 70.81% |
| SVM | 70.14% | 62.41% | 75.98% | 68.32% |
| MLP | 73.12% | 63.00% | 78.63% | **72.82%** |
| LightGBM | 72.43% | 61.13% | **80.74%** | 66.31% |
| Random Forest | 72.01% | 72.32% | 72.32% | 67.32% |
| Our Model | **76.06%** | **78.95%** | 74.89% | 65.44% |

**Table 4:** Comparison between our model and the deep learning models trained and validated by [17] for AD vs CN. The results are the weighted average of the performances of 10 cross-validation runs with random splits. The best value in each column is in bold.

| AD vs CN | Accuracy | Sensitivity | Specificity | F1 Score |
| --- | --- | --- | --- | --- |
| DeepConvNet | 54.21% | 45.43% | 57.59% | 47.01% |
| EEGNet [21] | 41.00% | 47.20% | 37.67% | 42.04% |
| EEGNetSSVEP [22] | 51.46% | 56.78% | 45.39% | 51.82% |
| DICE-net | **83.28%** | 79.81% | **87.94%** | **84.12%** |
| Our Model | 77.11% | **81.69%** | 72.49% | 78.17% |

**Table 5:** Comparison between our model and the deep learning models trained and validated by [17] for FTD vs CN. The results are the weighted average of the performances of 10 cross-validation runs with random splits. The best value in each column is in bold.

| FTD vs CN | Accuracy | Sensitivity | Specificity | F1 Score |
| --- | --- | --- | --- | --- |
| DeepConvNet | 64.21% | 62.41% | 37.05% | 60.20% |
| EEGNet [21] | 46.00% | 42.20% | 57.46% | 43.65% |
| EEGNetSSVEP [22] | 61.46% | 53.51% | 75.00% | 52.43% |
| DICE-net | **74.96%** | 60.62% | **78.63%** | **62.27%** |
| Our Model | 74.16% | **89.09%** | 70.69% | 56.49% |

Using LOSO validation, the model reaches an accuracy of 81.13% for AD vs CN and 76.06% for FTD vs CN. These are 4.1 points above the best baseline for AD (Random Forest, 77.01%) and 2.9 points above the best baseline for FTD (MLP, 73.12%). The sensitivity is also higher than for the best baseline, by 4.7 points for AD and 6.6 points for FTD. This comes with a lower specificity than the best baseline, by 2.1 points for AD and 5.9 points for FTD. The F1 score is the highest for AD vs CN but not for FTD vs CN, where our model has 65.44% and the MLP baseline has 72.82%.

Against the deep learning models in the random cross-validation setting, DICE-net has the highest accuracy for AD vs CN (83.28% versus 77.11% for our model), and the two models are close for FTD vs CN (74.96% versus 74.16%). Our model has the highest sensitivity for both tasks. DeepConvNet, EEGNet [21] and EEGNetSSVEP [22] have a considerably lower performance than our model on both tasks.

### 3.2 Node Saliency Maps

Fig. 2 shows the importance of the nodes for AD vs CN, and Fig. 3 shows it for FTD vs CN. The maps show the importance of each node relative to the minimum and maximum importance in the map. For AD vs CN, the Alpha (8–13 Hz), Theta (4–8 Hz) and Gamma (30–45 Hz) bands are the most important, and the electrodes *O*2 and *P* 4 are the most important nodes in these bands. The electrode *T* 4 is also important in the Alpha, Theta and Gamma bands. For FTD vs CN, the Alpha band is clearly the most important, with the highest values at *O*2 and *P* 4, and *T* 6, *T* 4 and *T* 3 are also highlighted, while the other bands have a much lower importance. Compared with AD vs CN, the frontal electrodes *Fp*1 and *Fp*2 have a higher importance for FTD vs CN.

**Figure 2:**
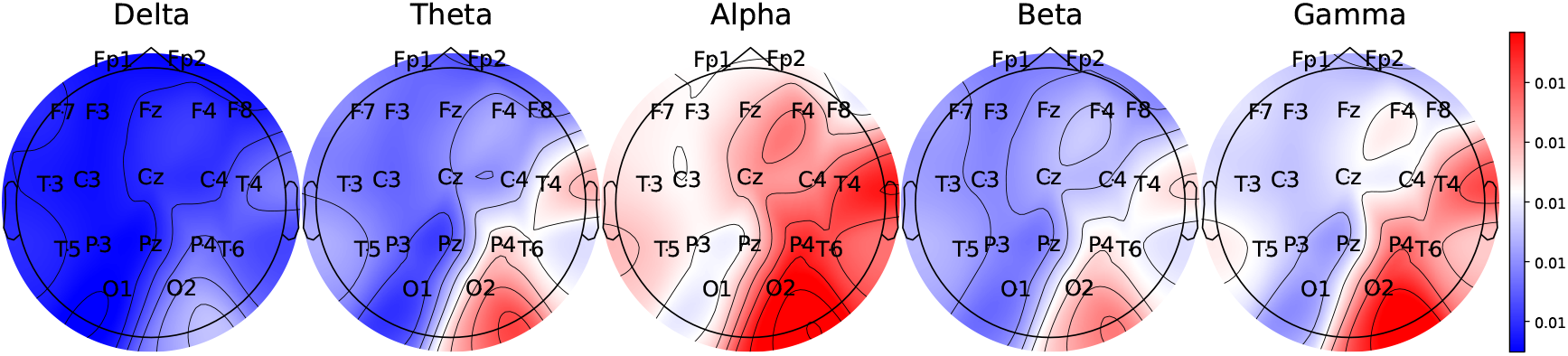
Importance of the nodes in the AD vs CN classification, using all node features and averaged over the LOSO subjects.

**Figure 3:**
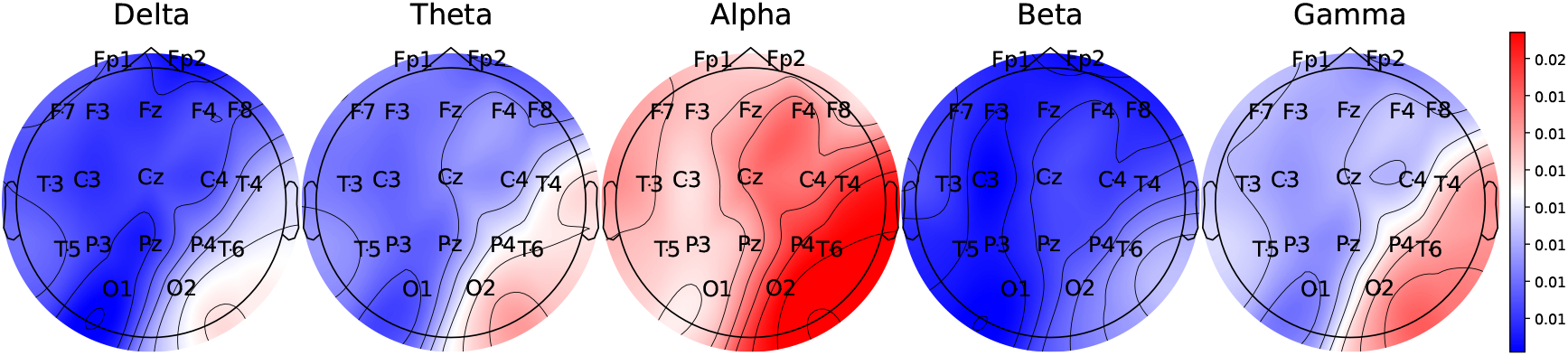
Importance of the nodes in the FTD vs CN classification, using all node features and averaged over the LOSO subjects.

### 3.3 Edge Saliency Maps

Fig. 4 shows the importance of the edges for AD vs CN. Like the node maps, the edge maps show the importance relative to the minimum and maximum values over all the edges. The Theta band is the most important band, followed by the Alpha and Delta bands, and the PLV edges have the highest importance, followed by Coherence. For FTD vs CN (Fig. 5), the Theta band is also the most important, followed by the Gamma and Alpha bands, and the PLV edges again have the highest importance, followed by Coherence. The edge maps of the two tasks are therefore similar.

**Figure 4:**
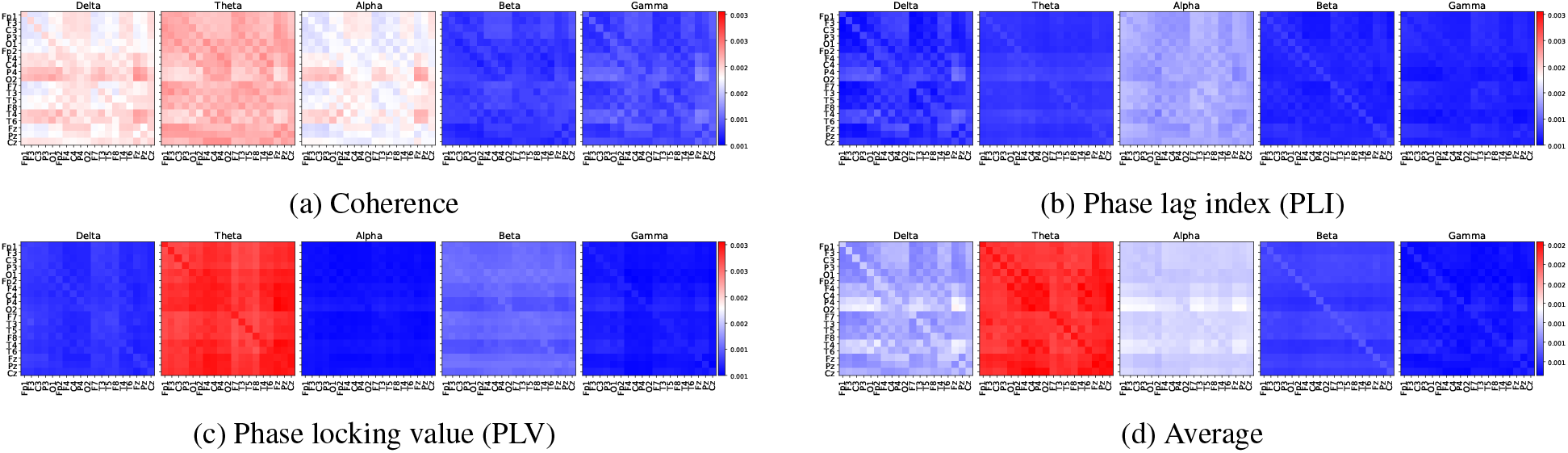
Relative importance of the edges for AD vs CN, for the different connectivity measures and frequency bands.

**Figure 5:**
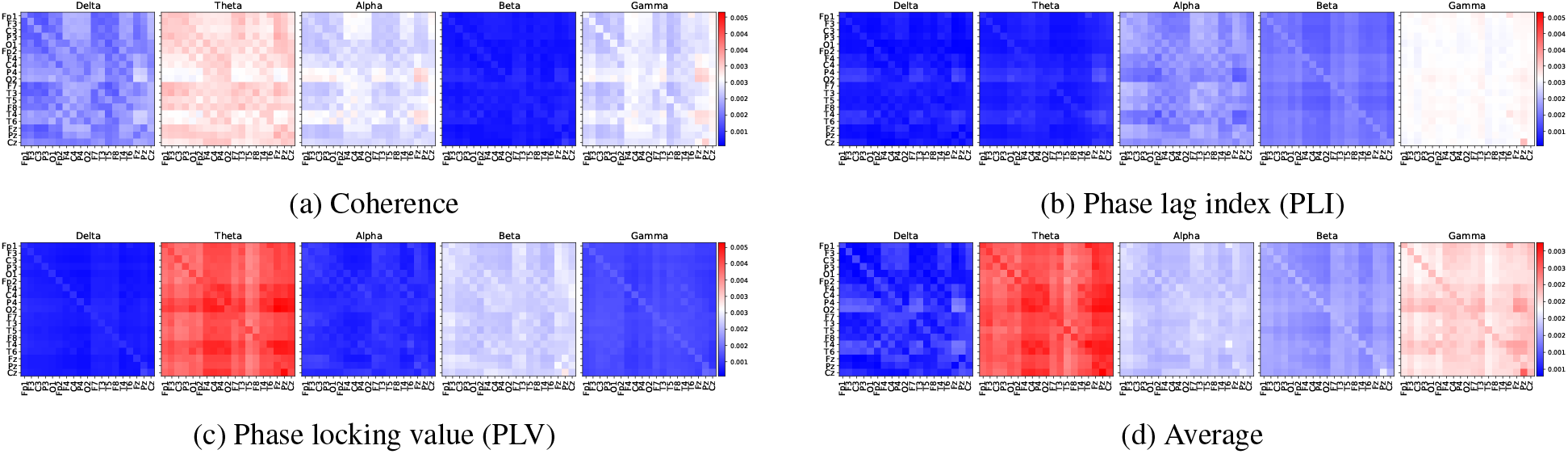
Relative importance of the edges for FTD vs CN, for the different connectivity measures and frequency bands.

### 3.4 Brain Network Maps

We look more closely at the networks that the edge maps highlight. For AD vs CN these are PLV in the Theta band and Coherence in the Theta, Alpha and Delta bands (Fig. 6). All three measures are symmetric, so the connections in these maps are symmetric by construction. In the Theta PLV network (Fig. 6a), the connections to and from *O*2, *O*1, *C*4, *P* 4 and *T* 4 are highlighted. This is in line with the node maps, where *O*2 and *P* 4 are the most important nodes. In the Alpha and Delta Coherence networks (Fig. 6c and 6d), the highlighted connections involve *O*2, *P* 4, *T* 4 and *T* 6 together with *Fz, Pz, Fp*1, *F* 3, *O*1 and *Fp*2, which belong to different brain regions. The connections between *Cz* and the other electrodes are also highlighted.

**Figure 6:**
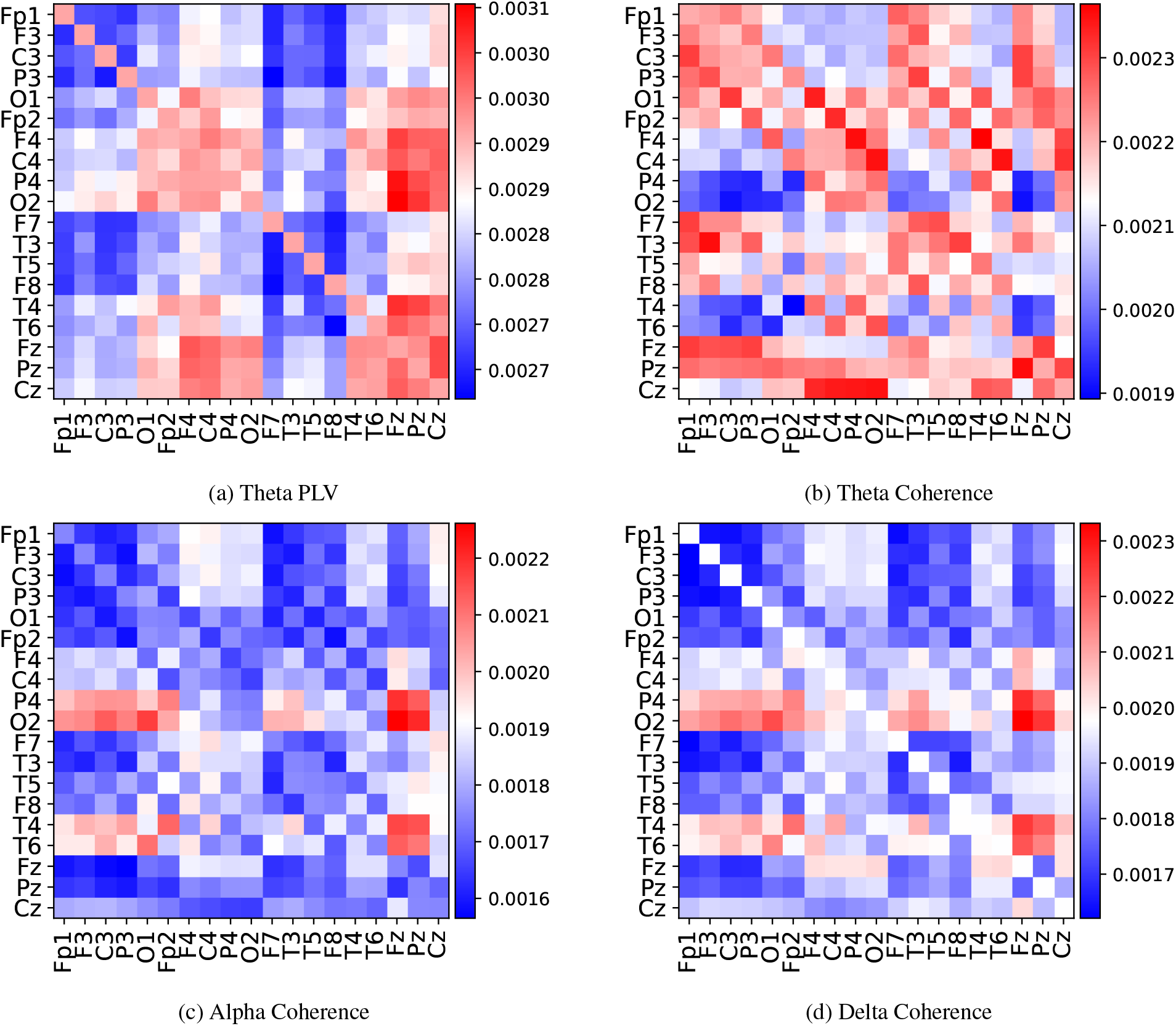
Relative importance of the edges for AD vs CN for the most important networks. Each plot is normalized to its own minimum and maximum to highlight the most important edges.

For FTD vs CN, the same networks are highlighted, and in addition the Gamma band networks of PLI and Coherence (Fig. 7). The Theta PLV network (Fig. 7a) is similar to the one for AD vs CN. The connections to and from *O*2, *C*4 and *P* 4 are highlighted, which is consistent with the node maps, where *O*2, *T* 6 and *P* 4 are the most important nodes. In the Coherence networks of the Alpha, Delta and Gamma bands (Fig. 7c, 7d and 7f), the highlighted connections involve *O*2, *P* 4, *T* 4 and *T* 6 together with *Fz, Pz, Fp*1, *F* 3, *O*1 and *Fp*2. The connections between *Cz* and the other electrodes are highlighted in Fig. 7a, 7b, 7c, 7d and 7f, and the connection between *Pz* and *Cz* is important in Fig. 7e and 7f.

**Figure 7:**
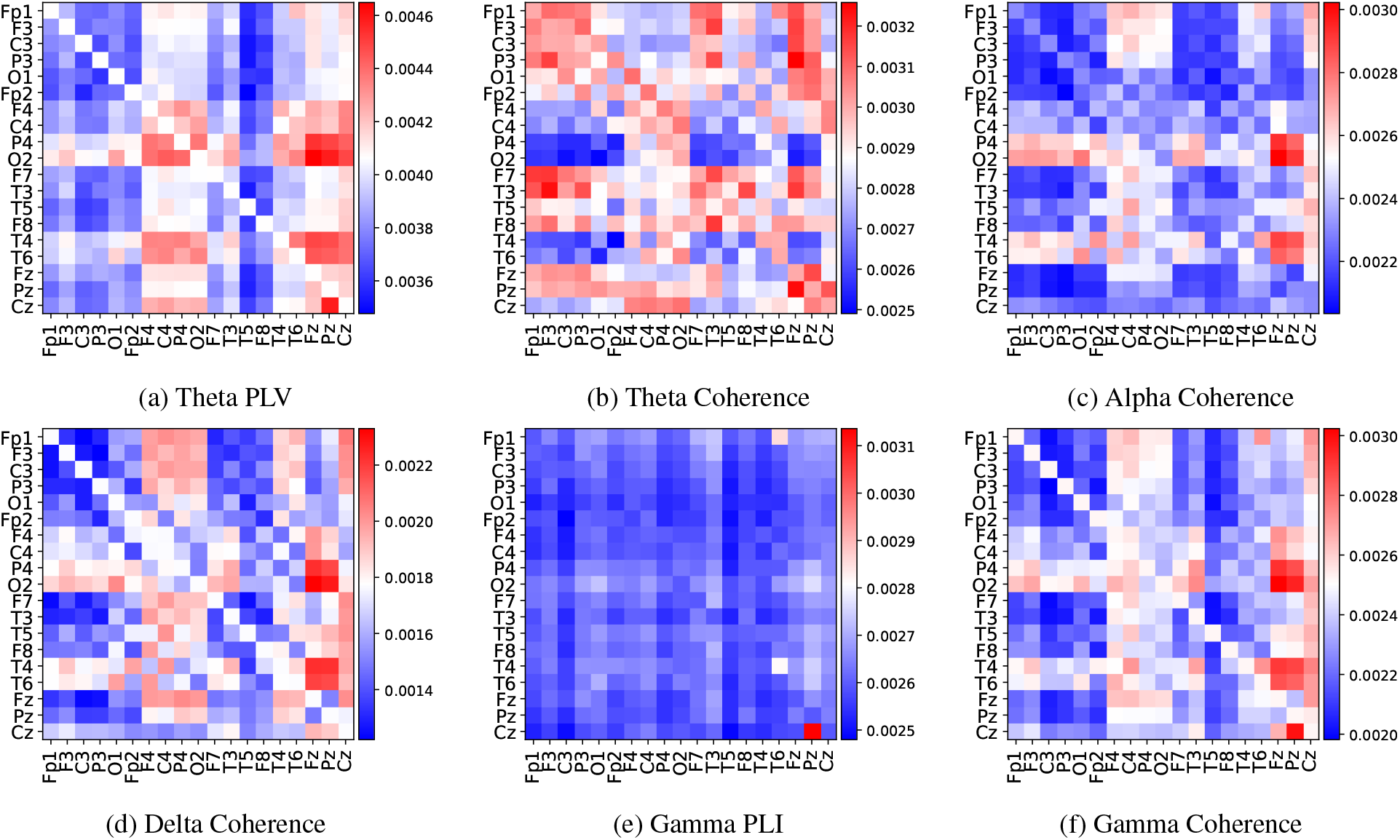
Relative importance of the edges for FTD vs CN for the most important networks. Each plot is normalized to its own minimum and maximum to highlight the most important edges.

### 3.5 Feature Ablation

To complement the saliency maps, we ran an ablation of the edge features on the AD vs CN task. In one set of variants, only one connectivity measure is used with its five frequency bands, so there are 5 edge features instead of 15. In another variant, the three measures are each averaged over the frequency bands, so there are 3 edge features, one per measure. The ablation was run for AD vs CN, which is the task that is new in this paper and has more subjects than FTD vs CN (65 versus 52).

Table 6 shows that Coherence is the best single measure in terms of accuracy (76.61%), while PLV and PLI are almost equal (73.72% and 73.59%). Using Coherence alone gives an accuracy 4.5 points below the full model, and averaging the edge features over the bands lowers the accuracy by 9.7 points. All of the ablated variants have an accuracy below the best classical baseline of Table 2 (77.01%). One subject corresponds to about 1.5 points of accuracy in this task, and the ablation was run once for each variant, so the smaller differences, such as the one between PLV and PLI, should not be interpreted.

**Table 6:** Feature ablation for AD vs CN. The results are the weighted average of the performances of the LOSO validation subjects. The number of edge features is 5 for each single measure, 3 for the averaged bands and 15 for the full model. The best value in each column is in bold.

| AD vs CN | Accuracy | Sensitivity | Specificity | F1 Score |
| --- | --- | --- | --- | --- |
| Coherence | 76.61% | 76.66% | 76.54% | 79.38% |
| PLI | 73.59% | 72.57% | 75.31% | 77.50% |
| PLV | 73.72% | 78.36% | 69.05% | 74.92% |
| Averaged bands | 71.41% | 72.99% | 69.33% | 74.35% |
| Full features | **81.13%** | **83.05%** | **78.85%** | **82.67%** |

## 4 Discussion

### 4.1 Classification Performance and Model Size

Under LOSO validation the model has a higher accuracy and sensitivity than the baselines reported with the dataset, on both tasks (Tables 2 and 3). The higher accuracy is accompanied by a higher sensitivity and a lower specificity than the best baseline. A high sensitivity is useful when the cost of missing a patient is high, but a lower specificity means more false alarms among the controls. The gain over the best baseline is 4.1 points for AD and 2.9 points for FTD, which corresponds to about three subjects for AD and one to two for FTD, and we have not tested whether the differences are larger than the variation between runs.

Against the deep learning models evaluated with random cross-validation (Tables 4 and 5), the accuracy of our model is close to DICE-net for FTD but about 6 points lower for AD, and it has a higher sensitivity on both tasks. The settings of the two groups of tables differ, so these comparisons are indicative only. The size of the models is very different. Our models have about 159 and 282 thousand parameters, while DICE-net has 170.5 million parameters according to [17]. This means that a competitive performance for FTD is reached with a model that is much smaller.

EEGNet and DeepConvNet, which use the raw EEG amplitudes, have a much lower performance than the models that use extracted features, which are ours and DICE-net, whose inputs are built from spectral coherence and relative band power. A possible explanation is that raw amplitudes are noisy and not comparable across recordings, because of differences in electrode placement and recording conditions. This is a hypothesis that we did not test, and the small size of the dataset may also play a role. In our own experiments, models trained on raw time series did not converge.

### 4.2 Frequency Bands

For the node features, the Alpha band is the most important in both tasks, followed by the Gamma, Theta or Beta bands. For the edge features, the Theta band is the most important, followed by the Alpha, Gamma or Delta bands, depending on the task. The importance of the Alpha band for node features is in line with the findings of [13] for Alzheimer’s disease. In the ablation, edge features averaged over the bands gave a lower accuracy than the band-resolved edge features, so keeping the bands separate helped in this experiment, although the number of edge features also differs. The ablation does not test the bands one by one.

The literature reports consistent abnormalities in the Delta, Theta and Alpha rhythms in Alzheimer’s disease [23, 24, 25, 26, 27, 28]. Relative Theta power is increased in patients with dementia due to AD [24], and global Theta power is increased in AD patients [23]. Other studies report an increase in Delta and Theta and a decrease in Alpha and Beta in AD patients [25, 26, 27, 28]. In patients with cognitive impairment, higher Theta and lower Beta relative power are reported in a longitudinal study [29]. For FTD, Alpha power is generally decreased, particularly in frontal and temporal regions [30], and lower Alpha activity was found in patients compared with controls in orbital frontal and temporal lobes [31]. FTD patients also show increased Theta power compared with controls [30, 32, 33]. Our saliency maps show a similar pattern, as Alpha and Theta are the most important bands, but saliency shows what the model uses and not the direction of the change in power, so we only claim that the bands match the literature.

### 4.3 Brain Regions

The occipital electrode *O*2 and temporal electrodes are the most important in the decisions of both classifiers. The occipital importance is strongest in the Alpha band, but it is present in nearly all bands. In addition, the frontal electrodes *Fp*1 and *Fp*2 have a higher importance in the FTD vs CN classifier. These results agree with those of [17], who analyzed the same data. The connectivity maps show the same regions as the node maps, with important connections to and from *O*2 and the temporal electrodes. A reduced occipital and frontal activity peak has been reported in mild Alzheimer’s disease compared with healthy individuals [34], and atrophy of the anterior temporal lobes is associated with semantic dementia [35, 36, 37, 38, 39]. Functional connectivity between the medial temporal lobe and the anterior-temporal system is reduced in the earliest stages of Alzheimer’s disease [40], and medial temporal lobe networks change with age and in MCI and are related to cognition and to later cognitive decline [41, 42]. These studies support the relevance of the regions that the model highlights, but scalp electrodes cannot be mapped directly to these deep or anterior structures, so the match is approximate.

### 4.4 Connectivity Measures and the Ablation

In the ablation, the full model with 15 edge features performs better than any model with a single connectivity measure or with band-averaged edge features. This is consistent with the idea that the different measures and bands carry information that the model can use. However, the variants also differ in the number of edge features (15, 5 and 3), so the ablation cannot separate the value of the information from the effect of the model input size. It was also run on the AD task only, so we do not know whether the same pattern holds for FTD, and once for each variant. We therefore treat it as a first indication and not as proof of complementary information.

The saliency maps and the ablation do not rank the measures in the same way. The saliency maps give the highest importance to PLV edges, followed by Coherence, while Coherence is the best single measure in the ablation. This is not a contradiction, because the two analyses ask different questions: saliency measures how sensitive the full model is to each input, and the ablation measures how informative one measure is on its own. It does mean that the saliency ranking should not be read as a ranking of how useful the measures are.

## 5 Conclusion and Limitations

We studied a GNN with frequency-resolved Coherence, PLV and PLI edge features for AD vs CN and FTD vs CN. Under LOSO validation it has a higher accuracy and sensitivity than the classical baselines reported with the dataset, and it is much smaller than DICE-net. Compared with DICE-net it has a comparable accuracy for FTD and a lower accuracy for AD, although the two were evaluated in a different setting. The saliency maps point to occipital and temporal regions, to the Alpha band for node features and to the Theta band for edge features, which agrees with earlier EEG studies. In the AD ablation, the full set of edge features performed better than a single measure or band-averaged edges, but the design does not separate information from model size.

The study has several limitations. The dataset is small (36 AD, 23 FTD and 29 control subjects), comes from one hospital, and has no external validation, and one subject changes the accuracy by about 1.5 to 1.9 points. The results are from single runs without confidence intervals or significance tests. The ablation was run on the AD task only, so we do not know whether the same pattern holds for FTD, and it does not control for the number of edge features. The saliency maps are compared with the literature, but we did not test their faithfulness, for example by removing the highlighted features. The input features are hand-crafted, and models on raw time series did not converge in our experiments, which we attribute to noise and the small dataset without testing it. The deep learning comparisons use results from another setting and not our own runs of those models. All windows are treated as independent samples, which ignores how the windows of one subject are related over time.

Future work could address these points by validating on an external dataset, running the ablation on both tasks with repeated runs and a control for the number of edge features, and testing the faithfulness of the attribution maps. Since Coherence, PLV and PLI are symmetric, directional measures such as Granger causality could be used to capture the direction of the coupling. Dynamic graphs could model how the windows of one subject are related, and cross-frequency features could be added as candidate biomarkers.

## Notes

### Competing Interest Statement

The authors have declared no competing interest.

